# Decoding yeast transcriptional regulation via a data-and mechanism-driven distributed large-scale network model

**DOI:** 10.1101/2025.02.08.636880

**Authors:** Xingcun Fan, Guangming Xiang, Wenbin Liao, Luchi Xiao, Siwei He, Hongzhong Lu, Xuefeng Yan

## Abstract

The complex transcriptional regulatory relationships among genes influence gene expression levels and play crucial roles in determining cellular phenotypes. Here, we present a novel distributed large-scale transcriptional regulatory neural network model (DLTRNM) by integrating prior knowledge into the reconstruction of pre-trained machine learning models followed with fine-tuning. Taken *S. cerevisiae* as the example, the transcriptional regulatory relationships, carefully compiled and documented, are utilized to define interactions between TFs and their TGs within DLTRNM. Subsequently, DLTRNM is pre-trained on pan-transcriptomic data and fine-tuned with time-series data, enabling it to accurately predict dynamic regulatory correlations between TFs and TGs, such as simulating TG responses by tuning the expression of TFs. In addition, DLTRNM can assist in identifying potential key TFs for a subset of genes, thus simplifying the complex and interrelated transcriptional regulatory networks (TRN). Based on the key TFs, it can also refine previously reported transcriptional regulatory subnetworks and highlight the core regulatory networks. DLTRNM offers a powerful tool for studying transcriptional regulation with reduced computational demands and enhanced interpretability. Thus, this study represents a significant step forward in systems biology for understanding the complicated transcriptional regulation within cells.

## Main

The transcriptional regulatory network (TRN) seeks to elucidate the reciprocal interactions among genes during transcription. These interactions result in variations in gene transcription rates, which subsequently affect gene expression levels, playing vital roles in essential biological functions such as cell differentiation, metabolism, immune response, stress response, and disease processes. The inference and modeling of TRNs have been thoroughly investigated to address the complexity and dynamics of TRNs under both genetic and environmental perturbations.

The structure of a TRN is typically characterized through Boolean, differential equation, and machine learning models. The Boolean model is the most intuitive, utilizing binary values (0 and 1) to represent the presence or absence of regulatory relationships between genes. In constructing the Boolean model, Boolean thresholds are employed to infer the most likely regulatory relationships based on gene expression data^1–5^. While binary networks require fewer parameters, they may fail to capture important details, such as regulatory information at low and moderate expression levels^6–8^. The second commonly used method involves the application of differential equations^9^. These models describe the dynamic changes in gene expression underlying transcriptional regulation^10–13^. The construction and parameterization of differential equations pose significant challenges, as each parameter within the equations requires prior knowledge for its definition^14^.

The continuous accumulation of omics data, such as transcriptomics, has further accelerated the inference of regulatory networks through the application of machine learning methods^15–17^. Several deep learning methods have also been employed in the construction of TRN^18^, such as CNN^19^, meta-learning^20^, GNN^21^. The application of deep learning in TRN construction has become widespread with the availability of large amounts of single-cell omics. Among these advancements, STGRNS is an interpretable method based on the Transformer architecture, specifically designed for inferring gene regulatory networks from single-cell transcriptomic data^22^. Further, scGREAT utilizes a Transformer backbone and biomedical language models, using spatial transcriptomics data as external validation, to uncover novel regulatory relationships between genes^23^. As the complexity of TRN models increases, pre-trained large models based on transfer learning have garnered increasing attention^24–27^ to probe gene regulation. However, until now, few pre-trained transcriptional regulation models have been developed for *S. cerevisiae*.

TRN models based on machine learning, particularly deep learning, are often regarded as black-box models, which makes their interpretation challenging^28^. Moreover, these machine learning-based procedures often fail to incorporate established transcriptional regulatory relationships. On the other hand, TRN models on top of fine-tuning large models also demand substantial computational resources, which are typically limited for most researchers.

To address the above issue, we propose a distributed large-scale transcriptional regulatory neural network model (DLTRNM) in order to synergistically integrate qualitative mechanistic knowledge with quantitative data-driven methods. Firstly, DLTRNM constructs a unique neural network-based subnetwork for each TG, which is built using transcriptional regulatory prior knowledge, forming a distributed structure that can operate independently. Next, the pretraining and fine-tuning transfer learning were employed to get the functional model. Afterwards, DLTRNM can simulate how TGs responses under in silico TFs perturbation, which, to some extent, could be verified using experimental dataset. It also facilitates the identification of key TFs within complex TRNs and inference of core regulatory networks. Lastly, DLTRNM could refine previously reported transcriptional regulatory subnetworks and advance the study of transcriptional regulatory networks. In summary, DLTRNM serves as a powerful computational tool to illustrate yeast metabolic regulation at a holistic level.

## Result

### Framework of DLTRNM

In this study, we propose a novel distributed large-scale transcriptional regulatory neural network model (DLTRNM) to reflect cellular transcriptional regulation at the system level. The architecture of DLTRNM incorporates three modules from the establishment of the network compositions to the model fine-tuning (Fig. 1).

**Fig. 1.**
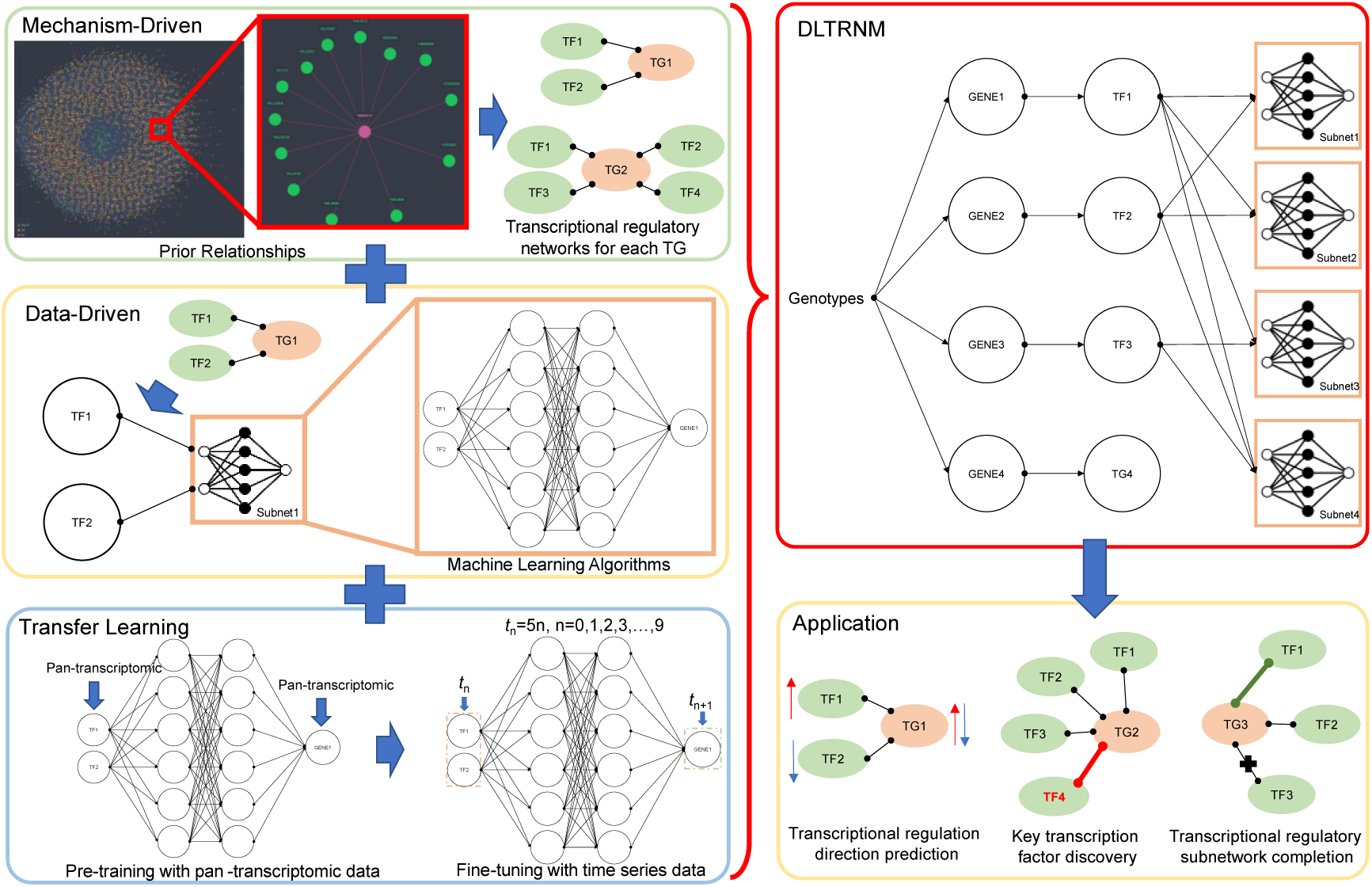
Overall framework of DLTRNM. DLTRNM integrates mechanism-driven modeling approaches with data-driven techniques, utilizing prior transcriptional regulatory relationships without directions sourced from databases as the foundational framework. It then employs machine learning methods, such as artificial neural networks, for data modeling. Additionally, DLTRNM leverages transfer learning to extract high-dimensional features from pan-transcriptomic data, which are subsequently applied to time-series prediction tasks, enabling the prediction of transcriptional regulation directions.

Firstly, prior knowledge of transcriptional regulation for *S. cerevisiae* was retrieved from databases, including SGD^29^ and YEASTRACT^30^, allowing us to elucidate the potential interactions between TFs and TG. These general interaction networks between TFs and TGs serve as one of the foundational elements for DLTRNM modeling. Next, a data-driven model was constructed using machine learning methods, such as artificial neural networks, with TG as the output and all TFs regulating the TG as inputs. Each TG is modeled independently, corresponding to a distinct subnetwork. The subnetworks are parallel and independent, ensuring that each subnetwork can be trained independently without interference from the others. Furthermore, DLTRNM incorporates transfer learning during the model training process. Initially, to capture the feature relationships between each TF and TG pair, the model is pre-trained using pan-transcriptomic data encompassing 969 high-quality transcriptomes derived from RNA sequencing of 1,032 natural isolates of *S. cerevisiae*^31^. This is followed by fine-tuning the model with time--course transcriptomics data collected in over 200 TF induction experiments^32^ to further learn how TFs dynamically regulated TGs.

### Training and fine-tuning of DLTRNM

During the pre-training phase of DLTRNM, each subnetwork was processed independently, enabling it to capture feature information from the pan-transcriptomic data. Based on the cumulative probability distribution of the loss function for the 6,085 subnetworks after pre-training in DLTRNM, 73.9% of the subnetworks had a sum of variances below 100 (Fig. 2A).

**Fig. 2.**
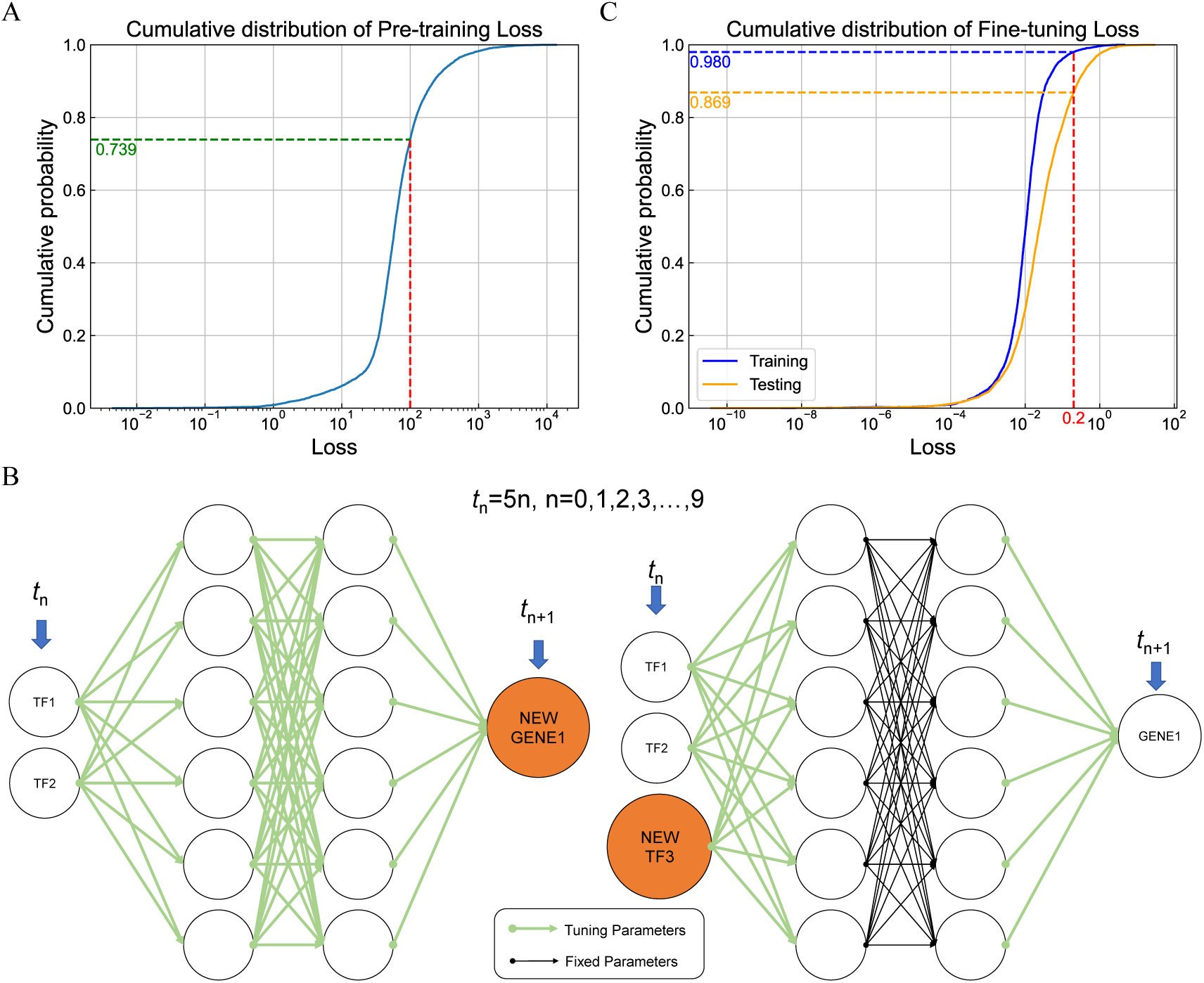
The pre-training and fine-tuning strategies and results of DLTRNM. (A) The cumulative probability distribution plot of the errors for 6,085 subnetworks during the pre-training phase. (B) Time series data has been organized sequentially to serve as input and output for fine-tuning each subnetwork in order to achieve transfer learning tasks. In cases where new TFs or TGs arise, the parameter training methods vary. The green line indicates parameters that require retraining, while the black line denotes fixed model parameters. (C) The cumulative probability distribution plots of training and testing errors for the fine-tuned 5,859 subnetworks on their respective datasets.

In the fine-tuning phase of the model, the original dataset (pan-transcriptomic data) was compared with the target dataset (time-series data), resulting in the identification of 5,859 genes as modeling targets for transfer learning. This included 5,855 genes common to both datasets, along with 4 newly added TFs (Methods). These results led to two scenarios during transfer training: the introduction of new TFs and new TG. For a new TG, since this gene does not exist in the original dataset, the entire network must be retrained from scratch. When a new TF appears for a specific TG, the new TF is introduced as a new input. The hidden layer weights of the original neural network are fixed, while others are retrained to complete the transfer learning process (Fig. 2B).

During the fine-tuning phase, DLTRNM benefited from aggressive parameter settings, which enabled it to better capture the characteristics of the time-series data. For each subnetwork, interpolation was applied to the time-series dataset to align the time intervals (Methods). Additionally, the interpolated dataset underwent quality filtering, removing data where gene expression levels remained unchanged or exhibited minimal variation. Such data were considered ineffective for reflecting the changes in the expression level of the TF-TG pairs. The results showed that 98.02% (5,743) of the subnetworks in DLTRNM achieved a training loss function value of less than 0.2 during the fine-tuning phase, with an average loss of 0.03. When testing the fine-tuned DLTRNM on the reserved test dataset, 86.87% (5,090) of the subnetworks had a loss of less than 0.2, with an average loss of 0.144 (Fig. 2C).

### Performances of DLTRNM in predicting dynamic correlations between TFs and its TGs

To assess the mode’s performance in dynamically predicting the time-series data after the fine-tuning of each subnetwork in DLTRNM, a semi-quantitative evaluation criterion has been established. Based on the difference between predicted values and actual values for each sample in the time-series dataset, the prediction performance of DLTRNM in each subnetwork at the sample level can be classified as “excellent” (high-precision predictions), “qualified” (directionally correct with tolerable errors), and “failed” (non-convergent or invalid results) (Fig. 3A, Methods). The output of the fine-tuned DLTRNM was firstly evaluated on the entire time-series dataset using above customized criteria (Fig. 3A). The results revealed that, on 82.84%, 3.41%, and 13.75% of the data samples, the performances of subnetwork models could be classified as “excellent”, “qualified”, and “failed”, respectively. This indicates that the majority of the predictions exhibited high accuracy for most samples.

**Fig. 3.**
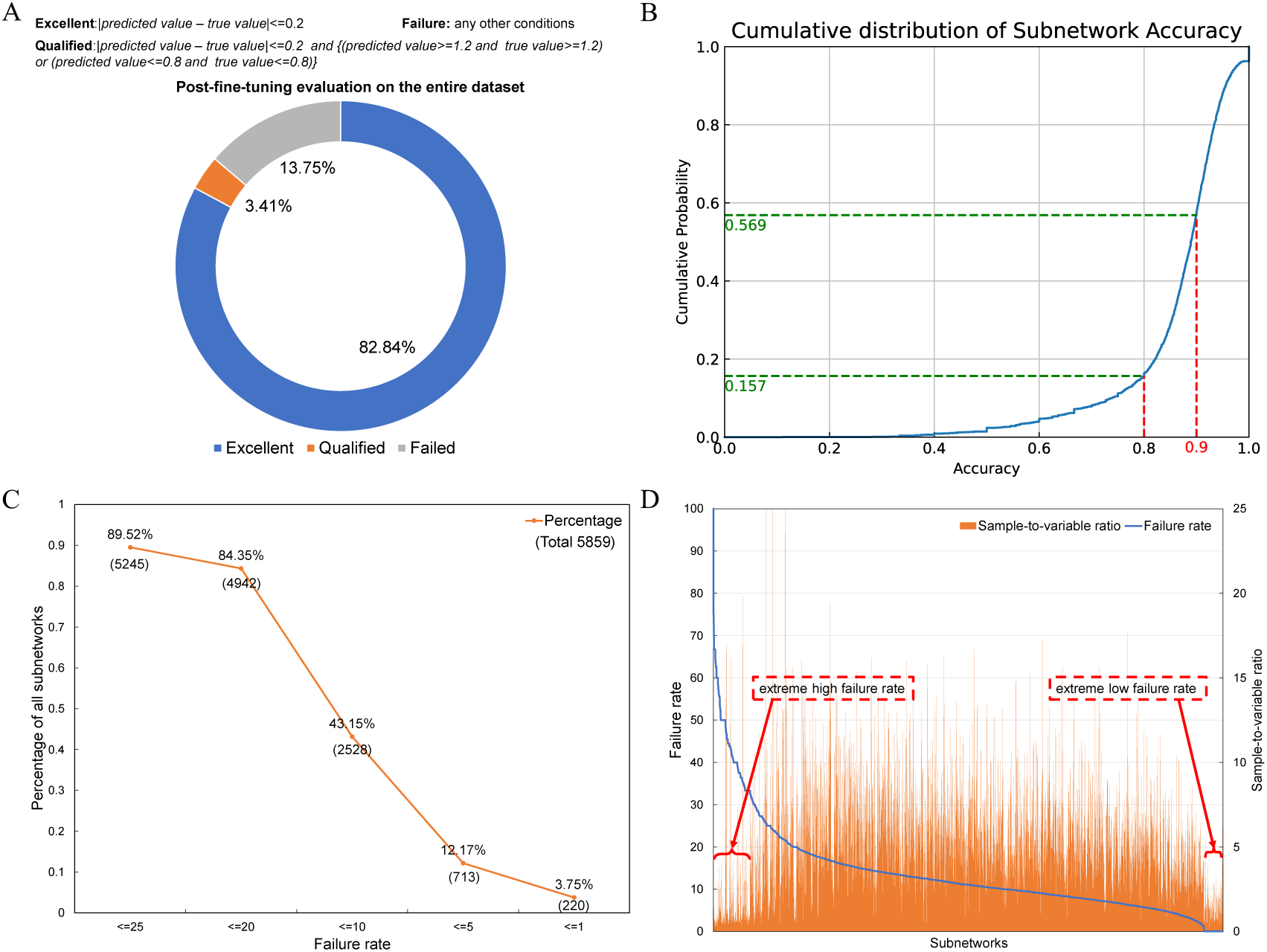
Results and analysis of fine-tuned DLTRNM based on custom evaluation criteria. (A) The custom evaluation criteria for predicted outputs, along with the overall model assessment across the entire dataset. (B) The cumulative probability distribution plot of accuracy for each subnetwork under the evaluation criteria shows that the majority of models exhibit accuracies ranging from 80% to 100%. (C) The statistical results of subnetworks filtered by varying prediction output failure probabilities indicate that 89.52% of the subnetworks achieve a failure rate lower than 25%. (D) The relationship between the amount of data and the number of input variables has a direct impact on the stability of the model.

To clearly represent the prediction accuracy of each subnetwork, the evaluation metrics “failure rate” and “accuracy” were defined based on the subnetwork-specific datasets. The “failure rate” was calculated as the percentage of samples classified as “failed” within each subnetwork’s datasets, while the “accuracy” was defined as the combined percentage of samples achieving either “excellent” or “qualified” predictions (Methods). 84.3% of the subnetworks achieved an accuracy of over 80%, with 43.1% of the subnetworks achieving an accuracy of over 90% (Fig. 3B). This indicates that the majority of the subnetworks exhibit high accuracy, while only a small number of subnetworks show significantly lower accuracy.

Furthermore, subnetworks were filtered based on different failure rate thresholds, and the number of subnetworks was statistically analyzed (Fig. 3C). In DLTRNM, 89.52% (5,245) of the subnetworks have a failure rate of less than 25%. As the accuracy criterion grows, the fraction and number of subnetworks that fulfill the threshold drop. When the failure rate is below 1%, only 3.75% (220) of the subnetworks meet this high-quality standard. Overall, the majority of subnetworks in DLTRNM exhibit good predictive performance, with a small portion demonstrating exceptional predictive capabilities.

It should be noted that the varying predictive performance of different subnetworks may be due to differences in the quantity and quality of their fine-tuning datasets. While the quality of the datasets is difficult to assess, the ratio between the number of datasets used to train each subnetwork and the dimensionality of its inputs can be analyzed to observe the impact of the dataset size on the performance of the subnetwork models (Fig. 3D). It can be observed that when the subnetwork’s failure rate is extreme, the aforementioned ratio is significantly smaller than that of other subnetworks. This suggests that when the training dataset size is too small relative to the number of model variables, the trained model becomes unstable. In the case of an excessively high failure rate, this could be due to insufficient data, where the model fails to adequately capture the underlying patterns, leading to poor generalization and a high failure rate. Conversely, when the failure rate is excessively low, it may suggest that the model has overfitted to the limited, homogenous data it was trained on, thus unable to generalize to other scenarios, and leading to an overly confident but inaccurate performance. Therefore, a well-balanced, high-quality, and sufficiently large dataset is crucial to generate accurate and stable subnetwork models.

### DLTRNM can independently simulate the how each TG was regulated in response to TF variations

After fine-tuning, DLTRNM, owing to its distributed structure, can independently simulate how the expression of each TG was induced by the variations of TFs (Methods), which accurately capturing the dynamic relationships between TFs and TGs without interference from other subnetworks.

To verify the performance of DLTRNM, the detailed regulatory relationship (i.e. positive or negative regulation of TGs by TFs) obtained from the SGD^29^, which was not taken account for model training, was used as the dataset for model validation. The regulation of TGs by TFs can be classified into two types: positive and negative, resulting in a total of 1,963 TF-TG relations. Here, each subnetwork in DLTRNM can be analyzed by up-regulating or down-regulating the corresponding TFs to simulated the tendencies in the expression of TGs under the perturbation of TFs. Due to the large number of relations, the complete results cannot be fully displayed, so a partial result is provided in Fig. 4A. It is found that, out of the 1,963 data points, 1,044 were accurately predicted, yielding an accuracy of approximately 53.2%. Notably, each subnetwork in DLTRNM is based on an artificial neural network structure, whose prediction results are likely to be nonlinear. As a result, when the input increases or decreases, the output may change in the same direction. In such instances, the model outputs were not unique, and the results are labeled as “other”.

**Fig. 4.**
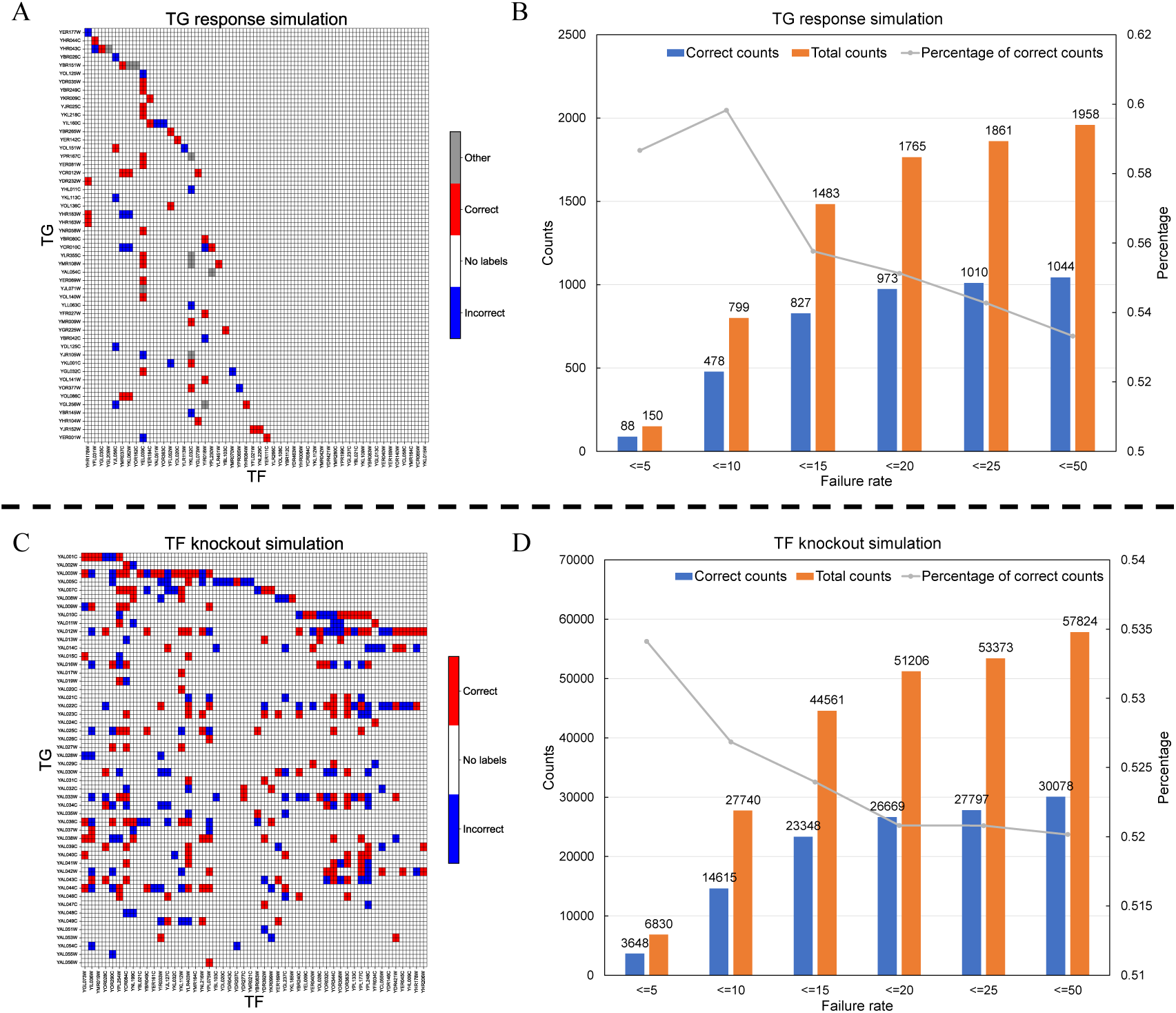
Results of DLTRNM in predicting TG response and simulating TF knockout. (A) A subset of heatmaps illustrates the variations of TFs corresponding to the predicted directional changes for each TG. Red squares indicate correct predictions across all aspects, blue squares represent incorrect predictions, and white squares signify the absence of labels for those conditions. Gray squares denote that the prediction of the TG, whether in the upregulation or downregulation direction of the TF, aligns with the label. (B) A statistical chart summarizes the total number of labels, correct predictions, and accuracy for all subnetworks that meet the failure rate criteria in TG response simulation. “Failure rate < 5” refers to subnetworks filtered based on the threshold of “failure rate < 5” in their evaluation results. Other labels correspond to results filtered using different thresholds. (C) A subset of heatmaps displays the predicted effects of knocking out each TF on all TGs. Red squares indicate correct predictions, blue squares represent incorrect predictions, and white squares signify the absence of labels for those conditions. (D) A statistical chart similarly demonstrates that the training results of the subnetworks directly influence the predictions of the impact of TF knockouts on TGs.

Due to variations in accuracy across subnetworks, the model’s prediction performances were statistically analyzed based on the subnetwork failure rate threshold (Fig. 4B). While “failure rate < 5” is expected to have higher accuracy, the observed accuracy was lower compared to “failure rate < 10”, likely due to insufficient data available for “failure rate < 5”. It is shown that other conditions, with more data points, followed the expected trend, highlighting the importance of having enough data to achieve reliable results.

### DLTRNM can systematically simulate the changes in expression of TGs after TF knockout

In addition to being analyzed as individual subnetworks, DLTRNM can also be applied systematically as a whole. DLTRNM enables the simulation of changes in all TGs resulting from the knockout of a specific TF. In this approach, the expression level of the targeted TF is set to zero, and a simulation is conducted across all subnetworks to observe the resulting changes in all TGs. The validation dataset used for simulating TF knockout was sourced from published literature^33^. The dataset includes the changes in expression of genes following TF knockout, which could indicate the potential correlation between TFs and TGs. If the expression trend in the TG after TF knockout aligns with the measured dataset, it is considered a correct prediction. Conversely, if the change does not align with the dataset, it is regarded as an incorrect prediction.

Knockout simulations were conducted for all 225 transcription factors associated with the 5,859 genes derived from the dataset in LDTRN. The results indicated that LDTRN accurately predicted 30,362 instances, yielding an accuracy of 51.95% (Fig. 4C). In addition, the results were statistically analyzed using the failure rate of each subnetwork as a threshold (Fig. 4D). Similarly, as the accuracy of the subnetworks increased, the accuracy of the predictions for the TF knockout data also improved.

### DLTRNM can predict key TFs determining the expression of TGs

Identifying key TFs for each TG is crucial for understanding the mechanisms underlying gene expression, especially in the context of designing engineered strains with specific traits. The distributed structure of DLTRNM subnetworks allows for the identification of TFs that have the most significant impact on TG expression. To pinpoint key TFs within TRNs, SHAP analysis was conducted on each subnetwork of DLTRNM (Methods). It should be noted that the TF with higher SHAP value is likely more critical in regulating TG expression. Thus, TFs with the highest SHAP values inferred by the model was classified as the key TFs for the corresponding TGs.

Based on the identification of key TFs, the TRN can be simplified by disregarding non-key TFs for each TG. Starting from the original TRN, the network was simplified by retaining the top 5, top 3, and top 1 key TFs (Fig. 5A, drawn by Cosmograph^34^). This simplification process reduced the original complex and coupled network into a clearer structure, thereby facilitating the identification of the core TRNs. Node degree analysis was performed on both the original and simplified networks. The simplified network exhibited a smaller node degree distribution, which was more conducive to revealing the core structure in reflecting how TFs regulate the expression of TGs (Fig. 5B).

**Fig. 5.**
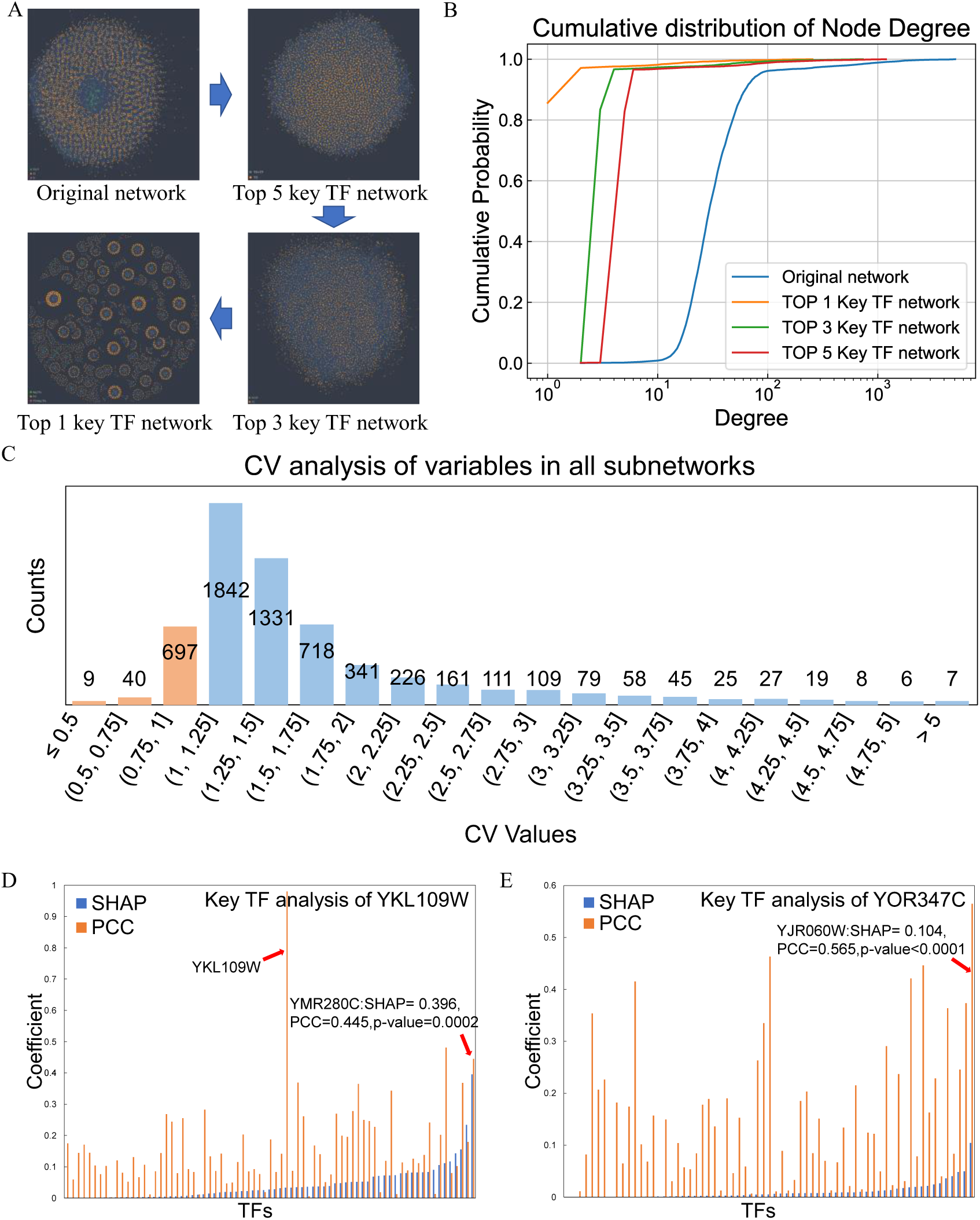
Results and analysis of key TF identification. (A) Utilizing the key TFs of each TG to simplify the overall TRN. (B) The cumulative probability distribution plots of the node degrees for the original relationship network and the simplified relationship network, which show a significant reduction in complexity. (C) Coefficient of Variation (CV) analysis of variables in all subnetworks. More than 87.27% of the subnetworks have large CV values, indicating significant differences in the importance of most variables across the networks. The SHAP, PCC, and p-values for the subnetworks of the target genes YKL109W (D) and YOR347C (E) are provided. The orange lines represent the SHAP and the blue ones represent the PCC.

Using the DLTRNM model, we successfully identified the key TFs that significantly contribute to the changes in expression of TGs. We further calculated the Coefficient of Variation (CV) of the SHAP values for each subnetwork to verify whether the calculated SHAP values effectively help identify key TFs (Fig. 5C) (Methods). Performing CV analysis on the SHAP values of TFs in each subnetwork aims to prevent scenarios where the SHAP value distribution within a subnetwork is overly concentrated, thereby ensuring that the identified key TFs exhibit significant importance to the model. In DLTRNM, 87.27% (5,113 out of 5,859) of the subnetworks had a CV value greater than 1. For these subnetworks, SHAP values exhibited distinct distribution patterns, allowing for clear identification of contribution of each TF to the model’s output. Overall, it indicates that DLTRNM can reflect the nonlinear or complex interactive relationships between TFs and TGs.

Further, we selected two TGs (YKL109W, which was associated with ethanol metabolism^35^, and YOR347C, which was associated with pyruvate metabolism^36^) for more detailed analysis. In addition to the SHAP values, we also calculated the Pearson Correlation Coefficients (PCC) between each TF and TG using their express levels from the fine-tuned dataset to further screen the key TFs inferred from the trained model (Fig. 5D and Fig. 5E) (Methods). Among these TFs, we selected variables with both high SHAP values and high PCC as potential key TFs. For YKL109W, the most influential TF was YMR280C, with a SHAP of 0.396 and a PCC of 0.445 (Fig. 5D, p-value = 0.0002). YMR280C is essential for the repression of multiple genes under non-fermentative growth conditions^37,38^, and may have a significant impact on YKL109W. For YOR347C, the most influential TF was YJR060W, with a SHAP of 0.104 and a PCC of 0.565 (Fig. 5E, p-value < 0.0001). YJR060W is directly involved in centromere function, and the deletion of this gene may lead to chromosome instability^39,40^. It is still not clear how YJR060W influencing the expression of YOR347C, thus the novel analysis here could provide clues for guiding the future experimental studies.

### DLTRNM can refine the existing transcriptional regulatory subnetworks

Lastly, we found that DLTRNM can refine previously proposed transcriptional regulatory subnetworks and identify key transcriptional regulatory interactions within these subnetworks.

The transcriptional subnetworks of the 12 most highly overexpressed *S. cerevisiae* genes in response to temperature changes were analyzed ^41^. DLTRNM encompasses almost all the regulatory relationships within this subnetwork and identifies potential key regulatory interactions based on key TF analysis (Fig. 6A). Interestingly, after reviewing both the SGD and YEARTACT databases, it was found that YBR193C is not listed as a TF. Based on the functional description of YBR193C, it is a subunit of the RNA polymerase II mediator complex and forms the RNA polymerase II holoenzyme by binding with the core polymerase subunits^42,43^. While it may be important for transcriptional regulation, its role may be not directly a TF. Therefore, the two transcriptional regulatory relationships involving YBR193C and YFR053C, as well as YBR193C and YCL040W could be refined through this subnetwork.

**Fig. 6.**
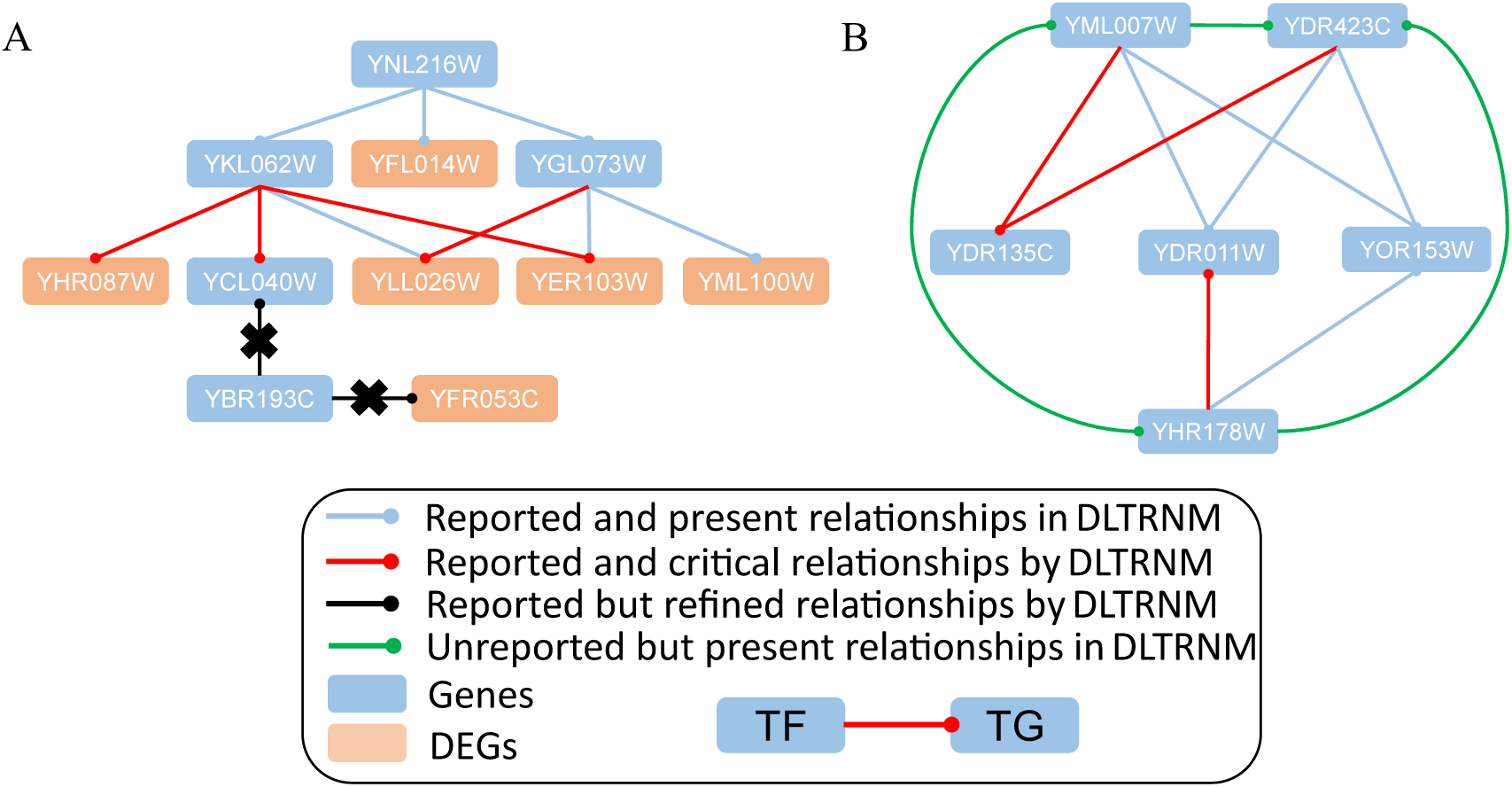
DLTRNM could refine the existing transcriptional regulatory subnetworks. (A) The regulatory network in *S. cerevisiae* under environmental temperature changes, where red and blue connections represent the networks reported in the literature. Each circular node, connected by edges, represents a regulated gene, with orange nodes indicating differentially expressed genes as proposed in the original literature. Specifically, the red edges indicate the key transcriptional relationships identified by DLTRNM. The black connections are those removed by our model in the subnetwork. (B) The regulatory network in *S. cerevisiae* under oxidative stress. The definitions of nodes and connections in the figure are the same as in (A). The green connections are those added by our model to refine the regulatory network.

As the second example, DLTRNM could refine regulatory network of *S. cerevisiae* under oxidative stress compared to that in the literature(Fig. 6B)^44^. In this subnetwork, DLTRNM not only included all the connections from the original network but also proposed potential key regulatory relationships (YML007W-YDR135C, YDR423CW-YDR135C, and YHR178W-YDR011W). In addition, we found that the extra relationships among YML007W, YDR423C, and YHR178W revealed by DLTRNM are also supported by literature^45–48^, whereas the original network did not indicate any transcriptional regulatory relationships among these three genes. The DLTRNM takes into account the potential relationships between them during model training, preventing the possible omission of regulatory relationships. Therefore, DLTRNM could, to some extent, help to refine the reported transcriptional subnetworks.

## Discussion

This study introduces a novel distributed large-scale transcriptional regulatory neural network model (DLTRNM) for *S. cerevisiae*, integrating mechanistic principles with data-driven approaches to infer genome-level transcriptional regulation. DLTRNM leverages prior knowledge of transcriptional regulatory relationships and employs machine learning to construct independent subnetworks for each target gene (TG), enabling precise modeling of gene expression dynamics. The model demonstrates high accuracy in predicting dynamic correlations between transcription factors (TFs) and TGs, simulating gene expression responses to TF variations, and identifying key TFs. DLTRNM can simulate both from a systemic perspective, considering the entire network, and from an independent perspective, focusing on individual subnetworks. DLTRNM also refines existing transcriptional regulatory subnetworks and highlights key regulatory interactions. By combining mechanistic insights with advanced machine learning techniques, DLTRNM offers a powerful tool for understanding and predicting transcriptional regulation, with applications in both independent and systemic analyses of gene regulatory networks.

However, DLTRNM heavily relies on the accuracy of the transcriptional regulatory relationships, as these directly influence the input variables of each subnetwork. If the prior knowledge is inaccurate, it will inevitably impact the simulation accuracy of DLTRNM. Moreover, discrepancies in the quality of the datasets for each subnetwork contribute to variations in performance. Previous studies have demonstrated that transcriptional regulatory relationships are not stable^49^. Therefore, when the dataset does not encompass the entire spectrum of transcriptional regulatory changes, the trained network is unable to model the dynamic transcriptional regulation accurately. As the mechanisms of transcriptional regulation remain incompletely understood, data-driven methods using neural networks can be continuously updated and refined as new regulatory mechanisms are uncovered. Therefore, as more accurate transcriptional regulatory relationships are discovered, the inputs of each subnetwork in DLTRNM can be further refined, enabling the construction of a more precise transcriptional regulatory network. Additionally, new training data can be continuously incorporated, aiming to provide a more comprehensive, dynamic, and accurately quantified transcriptional regulatory relationship.

In conclusion, DLTRNM represents a groundbreaking advancement in systems biology. Its distributed architecture and integration of mechanistic knowledge with data-driven learning offer unparalleled insights into transcriptional regulation. This model not only accelerates our understanding of yeast transcriptional regulation, but also paves the way for more efficient and comprehensive analysis of gene regulatory networks across diverse species on earth.

## Methods

### Acquisition of transcriptional regulatory relationships

The overall structure of DLTRNM is determined by the transcriptional regulatory relationships it acquires. Therefore, the quality of these relationships is critical to the performance of DLTRNM. For each subnetwork, the input consists of a set of all TFs involved in regulating the TG. Due to the powerful fitting ability of machine learning, DLTRNM can fit the entire process effectively, even when the input vector contains redundant variables. However, if the input variables are overly compact, some key variables might be overlooked, resulting in the loss of crucial information. Consequently, we aim to capture as many transcriptional regulatory relationships as possible to ensure that all potentially key variables are included in the model.

The YEASTRACT (Yeast Search for Transcriptional Regulators and Consensus Tracking) database is a comprehensive resource containing regulatory associations for over 206,000 *S. cerevisiae* genes, derived from more than 1,300 bibliographies. Each regulatory association in the database has been meticulously annotated by reviewing relevant references^30^. In YEASTRACT, each TF is associated with its binding site and two sets of TGs: the documented targets and the potential targets. The documented target set contains TGs that have been confirmed through literature reports, while the potential target set includes TGs that lack direct literature support but are predicted based on other criteria or experimental data.

While it is crucial to gather as many transcriptional regulatory relationships as possible, the documented transcriptional regulatory relationships are considered to be of higher quality and reliability. Including both the documented and potential target sets from YEASTRACT may introduce redundancy into the subnetwork, which could increase the complexity and computational cost of model construction and training. Therefore, DLTRNM uses only the documented TGs for each TF as the basis for transcriptional regulation relationships.

A statistical analysis of the transcriptional regulatory relationships revealed that most genes in *S. cerevisiae* are regulated by between 10 and 50 TFs, with a small subset of genes being regulated by more than 100 TFs (Fig. S1). This finding underscores the highly coupled and complex nature of the TRN in *S. cerevisiae*.

### Preprocessing of pan-transcriptome data preprocessing

In the pre-training phase of DLTRNM, the model was designed to acquire more generalized transcriptional regulatory relationships in *S. cerevisiae*. To achieve this, pan-transcriptomic data from 969 strains were utilized to pre-train DLTRNM with coarse precision. This pan-transcriptome dataset includes 969 high-quality transcripts derived from RNA sequencing of 1,032 natural isolates of *S. cerevisiae*^31^. The dataset contains at least 1 million mapped reads and is considered a high-quality representative of the *S. cerevisiae* pan-transcriptome. The dataset provides expression levels for 6,445 transcripts, including 4,977 core open reading frames (ORFs) and 1,468 auxiliary ORFs, in TPM format. To stabilize the pre-training process, the pan-transcriptome data were logarithmically transformed as described in equation (1).

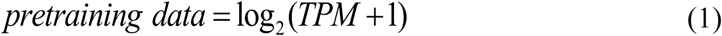

In this case, one was added to the original data to prevent zero values, which could otherwise result in vanishing gradients during model training.

### Preprocessing of time series data

After the pre-training phase, a fine-tuning method was employed to adapt DLTRNM for other prediction tasks. In this study, DLTRNM was designed to model changes in gene expression following the perturbation of each TF. The IDEA (Induction Dynamics Gene Expression Atlas) dataset is used to measure gene expression responses to hundreds of TF inductions^32^. The IDEA dataset consists of time-series data, with over 200 TF induction experiments, more than 20 million individual observations, and 100,000 dynamic responses that capture how genes in *S. cerevisiae* perform in response to TF perturbations. Since the IDEA data had already been log-transformed, they were further processed as described in equation (2) to facilitate transfer learning.

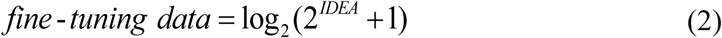

However, the time intervals in the IDEA dataset vary (Table 1).

**Table 1.**
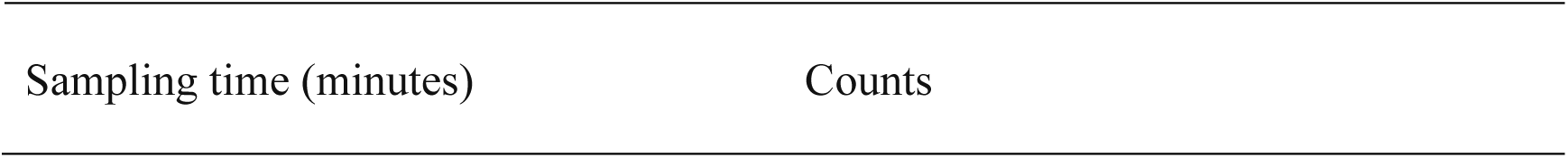
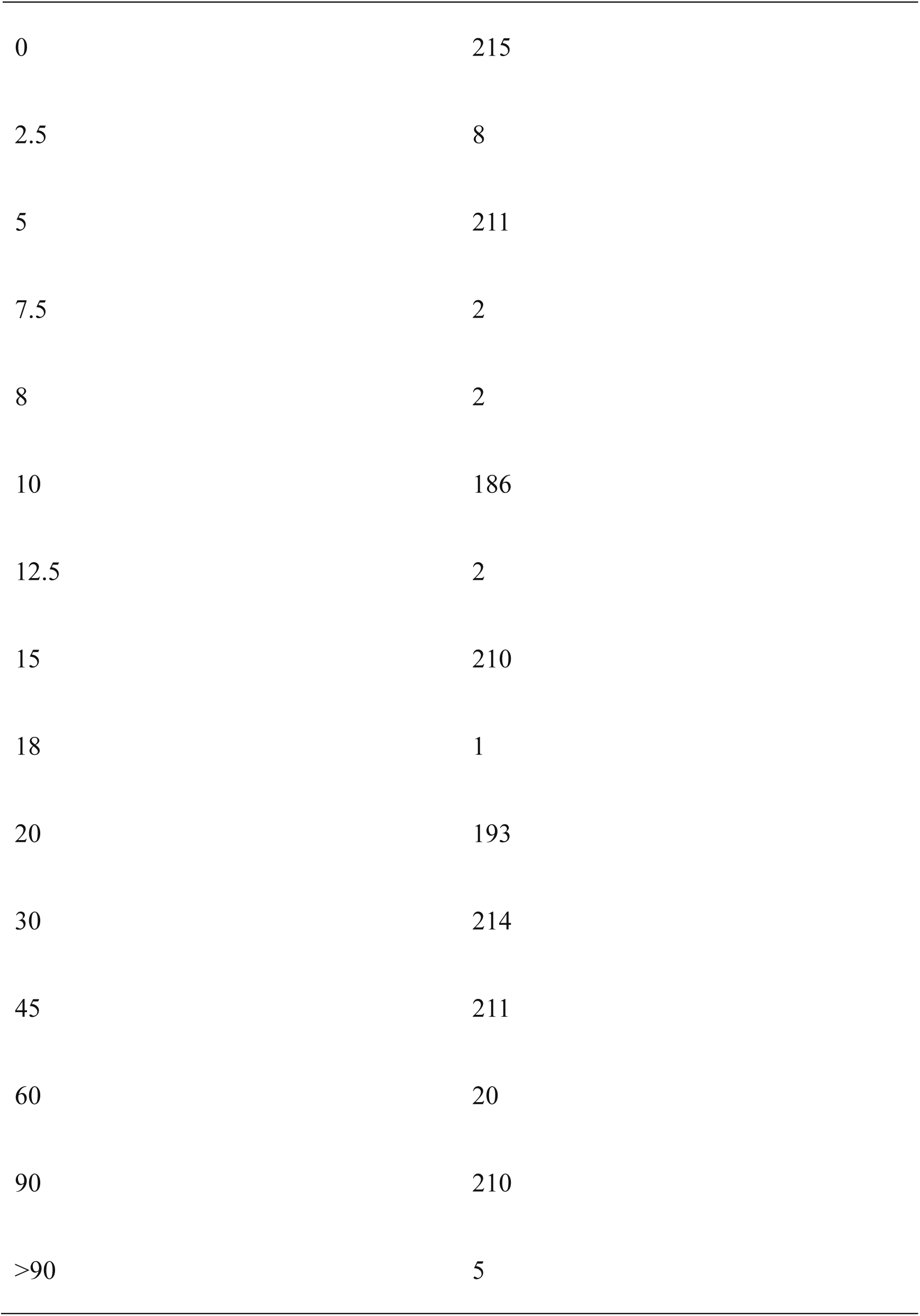
Sampling time points and total counts of time-series data in IDEA.

| Sampling time (minutes) | Counts |
| --- | --- |
| 0 | 215 |
| 2.5 | 8 |
| 5 | 211 |
| 7.5 | 2 |
| 8 | 2 |
| 10 | 186 |
| 12.5 | 2 |
| 15 | 210 |
| 18 | 1 |
| 20 | 193 |
| 30 | 214 |
| 45 | 211 |
| 60 | 20 |
| 90 | 210 |
| >90 | 5 |

It is evident that the IDEA dataset was sampled more frequently at specific time points, namely 0, 5, 10, 15, 20, 30, 45, and 90 minutes. Generally, uniform sampling intervals are preferred for time-series prediction tasks. Therefore, Radial basis function (RBF) interpolation, based on the Gaussian function in SciPy^50^, was used to complete the data. Given the high-frequency sampling times in IDEA and the challenge of accurately interpolating the long-time intervals from 45 to 90 minutes, the interpolation was focused only on the data within the first 45 minutes at 5-minute intervals. This process generated a time-series dataset spanning from 0 to 45 minutes, with 10 sampling points. An example of the interpolated data is shown (Fig. S2). This dataset was then applied to the fine-tuning phase of DLTRNM.

### Alignment of the two datasets

Since DLTRNM requires transfer learning between the aforementioned pan-transcriptomic data and time-series data, it is crucial to compare the differences between the two datasets, including the number of genes, TFs, and other relevant information (Fig. S3). The original dataset refers to the pan-transcriptomic dataset, while the target dataset refers to the time-series dataset. Upon comparison, it was found that the original and target datasets shared 5,855 common genes, with each dataset containing 230 unique genes, and the target dataset having 90 additional genes. Additionally, the target dataset contained four more TFs than the original dataset. Given that TFs are key variables in the regulatory network, they should be fully incorporated into the model whenever possible. The 221 TFs common to both datasets were included among the 5,855 shared genes. Therefore, a total of 5,859 genes were selected as the modeling targets for the transfer learning process.

### Distributed subnetworks and pre-training

The artificial neural network structure used for each subnetwork consists of two hidden layers. The number of nodes in the hidden layers is an empirically set hyperparameter, determined by the need to capture the three possible regulatory relationships of TFs on TGs: enhancement, repression, and no significant effect. To accommodate these relationships, the number of hidden layer nodes has been set to three times the number of input variables. This design ensures that the model can effectively capture and distinguish between these regulatory effects.

For example, in Fig. 1, the input of subnetwork 1 is a two-dimensional variable consisting of data from TF1 and TF2, corresponding to the TRN of TG1, which is regulated by these transcription factors. Subnetwork 1 is a fully connected neural network with a two-input layer, two hidden layers with six nodes each, and a single-output layer. This architecture is designed to capture the regulatory relationships between the TFs and TG1, enabling the model to predict the impact of these transcription factors on the target gene’s expression.

The entire dataset needs to be segmented for the distributed training of each subnetwork. The pre-training dataset is extracted from the paired data based on the TFs and TGs of each subnetwork, forming a unique dataset for each subnetwork. Since this was the pre-training phase, testing and validation were not required. Therefore, thxe entire reverse transcriptome dataset was used for training, enabling the model to better capture regulatory relationships. Additionally, an all-ones dataset, indicating no change in gene expression, was manually added to ensure the model could learn the baseline state of gene expression.

DLTRNM consists of nearly 6,000 subnetworks. Customizing unique hyperparameter settings for each subnetwork would be resource-intensive, so a unified set of hyperparameters was applied to all subnetworks. Since the pre-training phase involved coarser precision training, some hyperparameters were set more conservatively. In this study, the maximum epoch for pre-training was set to 2,500, the optimizer’s learning rate was 0.005, and the total variance loss function was used as presented in Equation (3).

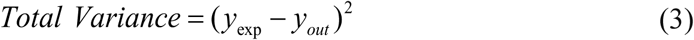

*y*_exp_ represents the training data, and *y_out_* represents the model’s output. The total variance loss function was chosen due to its simpler form, making it computationally efficient and more suitable for large-scale optimization problems. Moreover, the pre-training phase focused on minimizing overall error and ensuring that the network was adequately trained, without refining metrics.

### Transfer learning for fine tuning

DLTRNM undergoes pre-training to create a coarse accuracy model, which is then fine-tuned through transfer learning to develop a more precise model for other prediction tasks. Fine-tuning the model after pre-training is a standard transfer learning approach, where some parameters are fixed while others continue to be trained. In DLTRNM, the weights between the two hidden layers of each subnetwork are fixed, while the weights of the input and output layers are fine-tuned based on the new dataset. As with pre-training, the interpolated time-series dataset for fine-tuning is segmented for each subnetwork. For each subnetwork, the TF data from the previous sampling point is used as input, while the TG data from the subsequent sampling point is used as output, simulating the response of the TG to changes in the TF.

A large number of TGs were found to have a value of 1 at sampling time points in the deflated time-series data, indicating that these data do not capture changes in TGs. This may be due to time lags in the response. As the amount of such data increases, the fine-tuned network’s ability to simulate changes in TGs could be significantly affected. Therefore, we performed data screening for the fine-tuning process. Data with minimal changes in TFs (0.8 < x < 1.2) or no change in TGs (y = 1) were first excluded from the fine-tuning datasets of each subnetwork. In other words, only data with significant changes in TFs and TGs were retained. Additionally, it was considered that data with minimal changes in TFs and no change in TGs reflect the inherent robustness of the regulatory process. To avoid impacting the modeling of TG changes, this data was set to represent 20% of the TG change data.

After the dataset was split into training and test sets with a 9:1 ratio, it was observed that the amount of training data for the minutiae network was very small. For subnetworks with insufficient data, using ten-fold cross-validation may consume excessive computational resources. Therefore, a hierarchical cross-validation strategy was applied to reduce computational resource consumption by varying the number of cross-validation repetitions based on the data volume of each subnetwork. Specifically, the cross-validation for subnetworks follows these rules: for datasets with fewer than 20 samples, training is repeated 5 times; for datasets with 20 to 30 samples, it is repeated 4 times; for datasets with 31 to 40 samples, it is repeated 3 times; for datasets with 41 to 50 samples, it is repeated twice; and for datasets with more than 50 samples, training is performed only once. The network with the lowest test error is then selected and saved.

In the fine-tuning phase, accuracy became a key requirement for the subnetworks, so the hyperparameters were set more aggressively and uniformly. The maximum number of epochs for each subnetwork was determined based on the size of its dataset, as presented in Equation (4). The upper bound for the number of epochs across all subnetworks was set to 20,000.

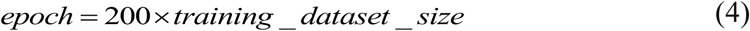

The optimizer’s learning rate was set to 0.001 with a weight decay of 0.001, and the mean squared error loss function was used, as presented in Equation (5).

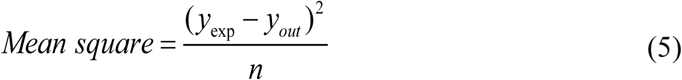

*n* represents the number of data points. *y*_exp_ represents the training data, and *y_out_* represents the model’s output.

### Custom validation criteria

The complexity and variability of transcriptional regulatory relationships pose significant challenges to the quantitative prediction of gene expression. Therefore, this study focused on semi-quantitative or qualitative changes in gene expression for model validation after fine-tuning. Evaluation criteria for each predicted result were developed based on the distribution of the output data from DLTRNM. The customized evaluation criteria were as follows.

When the absolute error between the predicted result and the labeled value was less than or equal to 0.2, the result was considered to exhibit high precision in numerical prediction, accurately simulating the quantitative transcriptional and regulatory relationships. The prediction result was classified as “excellent”.

When the absolute error between the predicted result and the labeled value was greater than 0.2, the prediction result was considered inaccurate in terms of quantitative indicators. However, if both values were simultaneously greater than 1.2 or less than 0.8, the trend of change was considered to be correctly predicted. Therefore, although the prediction in this case was not sufficiently accurate, it could still be considered correct and classified as “qualified”.

When the relationship between the predicted result and the labeled value did not fall into the categories above, the result was considered incorrect due to failing to meet the specified accuracy requirements in either numerical prediction or trend prediction. The prediction result was then classified as “failed”.

All data from each subnetwork could be evaluated according to this rule, and the accuracy of all networks could ultimately be visualized.

### Failure rate

The “failure rate” was used to evaluate the accuracy of the fine-tuned model on the dataset. After evaluating each data point based on the custom validation criteria, the results were classified into three categories: “excellent”, “qualified”, and “failed”. The calculation of the “failure rate” is defined by Equation (6) as follows:

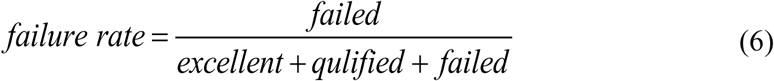

The “failure rate” provided an intuitive measure of the subnetwork’s performance following fine-tuning. A lower “failure rate” indicated higher accuracy of the subnetwork after fine-tuning, while a higher failure rate indicated lower accuracy. Thus, the “accuracy” can be defined by Equation (7) as follows:

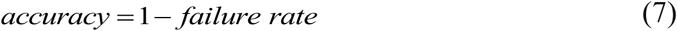

### TG response simulation

In DLTRNM, each subnetwork can be analyzed by modifying its input (i.e., TFs) by factors ranging from 2 to 5 times and from 0.2 to 0.5 times, in order to observe the resulting changes in its output (i.e., the TG). If the trend of change in TFs matches the trend of change in the TG (i.e., both increase or decrease simultaneously), it will be considered a positive correlation. Conversely, if the trends are opposite (i.e., when one increases while the other decreases), it will be considered a negative correlation.

### SHAP value

SHAP (SHapley Additive exPlanations) values were used to identify which input variable in each subnetwork of the fine-tuned DLTRNM had the greatest influence on the output, specifically to determine which TF most significantly impacted the expression change of the TG. SHAP is a method for interpreting the predictions of machine learning (ML) models, grounded in the concept of Shapley values from cooperative game theory. It calculates the marginal contribution of each feature across different feature combinations to assess its contribution to the model’s predictions. This approach provides a consistent and interpretable method for understanding complex black-box models and identifying the importance of individual features in the final prediction outcome. In this study, the SHAP package^51^ in Python was employed to compute and analyze SHAP values. The contribution of each variable was explained using the DeepExplainer on the fine-tuned dataset.

### Coefficient of Variation

The coefficient of variation (CV) is a statistical measure used to quantify the relative variability within a dataset, defined as the ratio of the standard deviation to the mean.

Mathematically, the CV is calculated as shown in Equation (8).

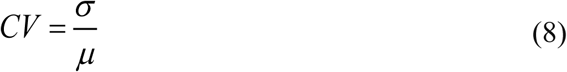

*σ* represents the standard deviation and *μ* denotes the mean of the dataset. This dimensionless quantity allows for the comparison of variability across datasets with differing units or scales, providing a normalized measure of dispersion. The CV is instrumental in assessing the consistency and stability of data, independent of the absolute magnitude of the measured variable. A higher CV signifies greater relative variability, whereas a lower CV reflects a more stable or consistent dataset. Consequently, the CV is a critical tool for comparing the precision of measurements, evaluating experimental reproducibility, and assessing data reliability across different conditions or populations.

In this study, the CV was employed to assess the variability in the importance of variables across subnetworks. A higher CV value indicated a greater ability to identify key variables within the subnetwork that are critical to the model.

### Pearson correlation coefficient

The Pearson Correlation Coefficient (PCC) was used to calculate the correlation between variables in the fine-tuned dataset, in order to verify whether the model has effectively learned the features in the data. The PCC is a statistical measure that quantifies the linear relationship between two variables. Its value ranges from −1 to 1, where −1 indicates a perfect negative linear relationship, 1 indicates a perfect positive linear relationship, and 0 signifies no linear relationship. The formula for the Pearson Correlation Coefficient is presented in Equation (9).

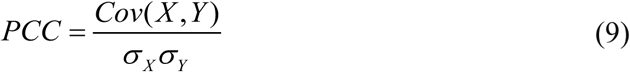

*Cov*(*X*, *Y*) is the covariance of variables X and Y. *σ _X_*, *σ_Y_* is the standard deviations of variables X and Y. The calculation of correlation coefficients is implemented using SciPy^50^. It should be noted that the PCC reflects only the linear relationship between two variables, whereas transcriptional regulatory relationships are often multivariate and nonlinear in nature.

## Date and code availability

All relevant data, codes and models are available in the GitHub: https://github.com/DCMO-ecust/DLTRNM.

## Acknowledgements

This work was financially supported by the National key research and development program of China (2020YFA0908300) and Shanghai Municipal Science and Technology Major Project.

## Declaration of generative AI and Ai-assisted technologies in the writing process

In the preparation of this manuscript, the authors utilized Large Language Models to enhance the clarity and flow of the text, as well as to refine the logical structure of the writing. All AI-generated content was rigorously reviewed, edited, and verified by the authors, who take full responsibility for the accuracy and integrity of the final work.

