## Supplemental figures for "Decoding yeast transcriptional regulation via a data-and mechanism-driven distributed large-scale network model"

### Supplementary information

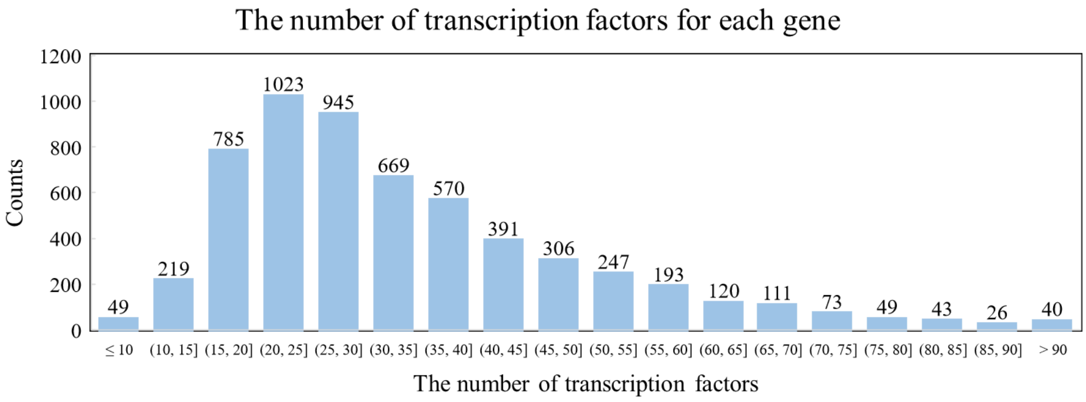


**Fig. S1 The number of TFs regulating each gene**. The majority of genes are regulated by dozens of TFs, indicating the complexity of the entire regulatory network.

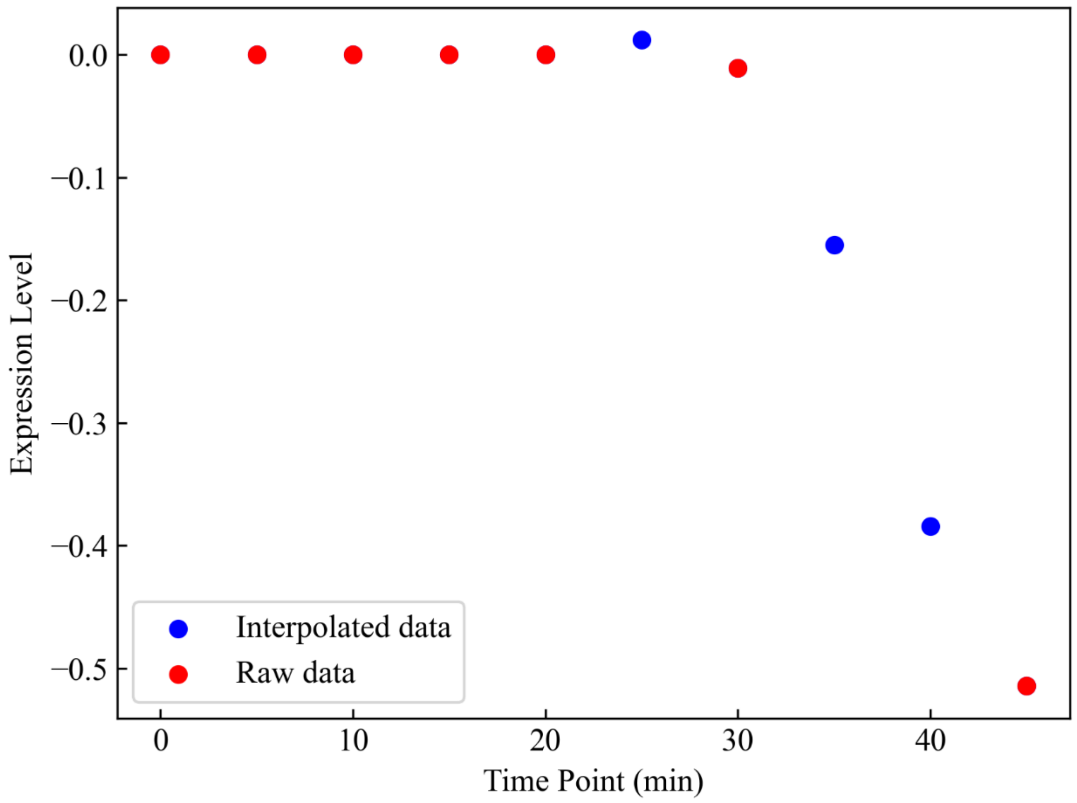


**Fig. S2 Example plot of interpolation for a time series dataset**. The red points represent the original data and the blue points represent the interpolated data.

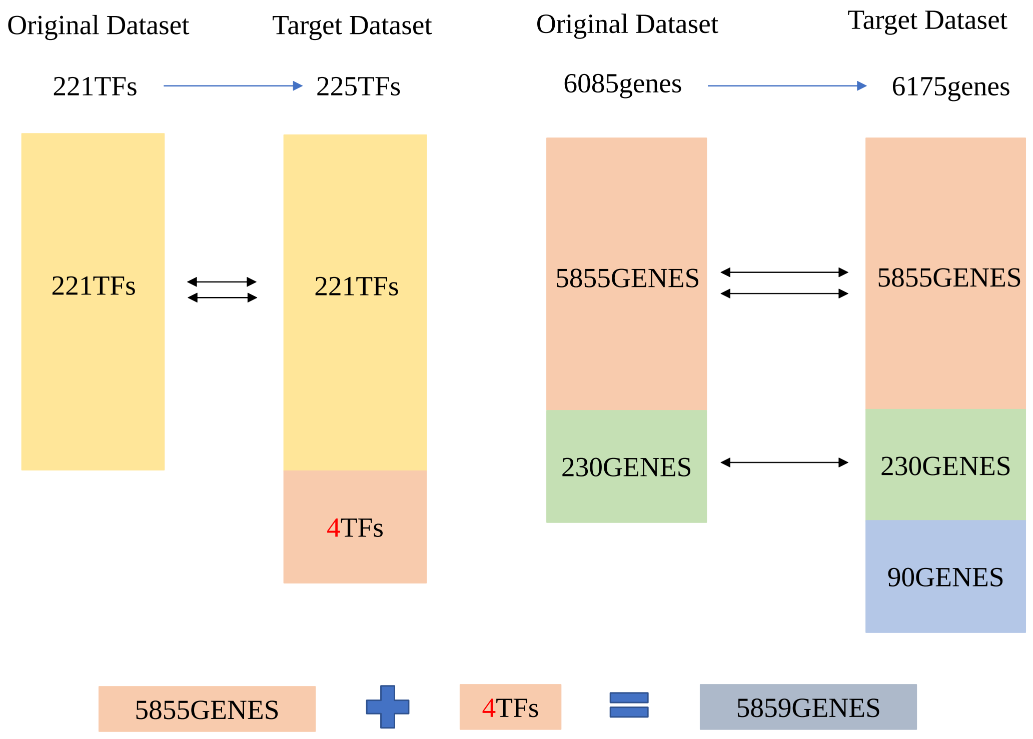

**Fig. S3 The comparison of the two datasets**. In transfer learning, the common 5855 genes (including 221 TFs) were selected, along with 4 additional TFs, as the new modeling targets.
